# The role of feedback in the production of skilled finger sequences

**DOI:** 10.1101/2021.07.02.450916

**Authors:** Nicola J. Popp, Carlos R. Hernandez-Castillo, Paul L. Gribble, Jörn Diedrichsen

**Affiliations:** The Brain and Mind Institute, University of Western Ontario, Canada; Faculty of Computer Science, Dalhousie University, Canada; Department of Psychology, University of Western Ontario, Canada; Department of Physiology & Pharmacology, University of Western Ontario, Canada; Haskins Laboratories, USA; Department of Statistical and Actuarial Sciences, University of Western Ontario, Canada; Department of Computer Science, University of Western Ontario, Canada

## Abstract

Actions involving fine control of the hand, for example grasping an object, rely heavily on sensory information from the fingertips. While the integration of feedback during execution of individual movements is well understood, less is known about the use of sensory feedback in the control of skilled movement sequences. To address this gap, we trained participants to produce sequences of finger movements on a keyboard-like device over a four-day training period. Participants received haptic, visual, and auditory feedback indicating the occurrence of each finger press. We then either transiently delayed or advanced the feedback for a single press by a small amount of time (30 or 60 ms). We observed that participants rapidly adjusted their ongoing finger press by either accelerating or prolonging the ongoing press, in accordance with the direction of the perturbation. Furthermore, we could show that this rapid behavioural modulation was driven by haptic feedback. While these feedback-driven adjustments reduced in size with practice, they were still clearly present at the end of training. In contrast to the directionally-specific effect we observed on the perturbed press, a feedback perturbation resulted in a delayed onset of the subsequent presses irrespective of perturbation direction or feedback modality. This observation is consistent with a hierarchical organization of skilled movement sequences, with different levels reacting distinctly to sensory perturbations.

## Introduction

Most motor behaviours strongly depend on feedback. When we grasp a full cup and feel a sudden slip, we can swiftly adjust our grip force to avoid the cup slipping from our hand. This correction can occur in less than 100 ms (Cole and Abbs 1988; Hernandez-Castillo et al. 2020; Johansson et al. 1992). Feedback from other senses such as vision (Day and Lyon 2000; Veerman et al. 2008) and audition (Burnett et al. 1998; Howell 2004) is also used for the control of an ongoing movements, albeit at slightly slower speeds (at 90-260 ms and 100-200 ms respectively). Based on the importance of sensory feedback, researchers have proposed that continuous feedback integration is essential for accurate movement execution (Adams 1971).

While much is known about the rapid sensory feedback integration during the execution of individual movements (for reviews see Cluff, Crevecoeur, & Scott, 2015; Scott, 2012; Shadmehr, Smith, & Krakauer, 2010), less is known about the integration of sensory feedback during the execution of sequences of finger movements. Previous studies investigating this topic have primarily focused on tasks in which participants were asked to synchronize their movements with an external pacing tone (Aschersleben 2002; Gates et al. 1974; Kulpa and Pfordresher 2013; Pfordresher and Benitez 2007; Repp 2000; van der Steen et al. 2014). Studies investigating the role of sensory feedback in tasks in which participants execute finger movements as fast as possible, however, are scarce (Jay and Hubbold 2005; Long 1975). Moreover, the majority of studies investigating this topic have focused on perturbing the slower visual or auditory feedback channels. Hence, these studies were unable to examine the full range of rapid feedback adjustments that are possible during a finger press.

Here we probed the use of sensory feedback during the execution of fast finger movement sequences. We manipulated haptic, visual, and auditory feedback on a few selected presses within a sequence, in a way that was not consciously perceivable by the vast majority of participants. Participants were trained on sequences of finger movements on an isometric keyboard throughout a four-day training period. On each press, upon reaching a given force threshold, participants were given a small haptic stimulus, similar to the feedback devices embedded in modern computer trackpads or smartphones. Concurrently, auditory and visual feedback indicated the successful pressing of the key. We then either delayed or advanced feedback on a single press within a sequence to probe how this sensory feedback is used in control. During the delayed feedback perturbation participants were not required to wait for the feedback to perform the subsequent presses – thus, by design, they could perform the task without considering feedback. However, we found an immediate, directionally-specific reaction to the feedback perturbation, providing strong evidence for the reliance of fast finger sequences on feedback.

The way participants react to a small feedback perturbation also provides a probe into how skilled motor sequences are organized. Models of sequence performance usually fall on a continuum along two extremes (Diedrichsen and Kornysheva 2015). On one side, sequences are controlled as a single unit or motor program (Keele 1968) that specifies the detailed muscle commands necessary to produce the sequence (Fig. 1a). On the other end is the idea that movement sequences are controlled hierarchically (Rosenbaum et al. 1983), in which one more abstract layer represents the sequence to be executed and a lower-level layer generates the detailed muscle commands for each finger press (Fig. 1b).

**Figure 1.**
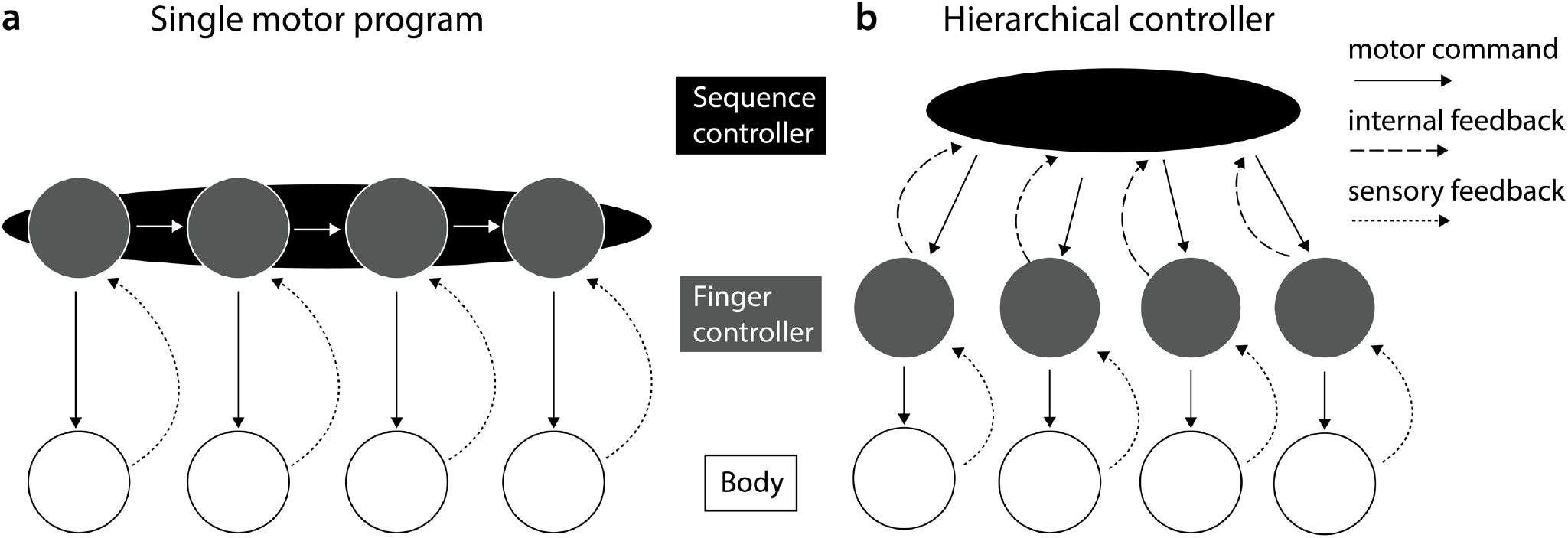
Two hypothetical representations of movement sequences. **(a)** A single motor program represents the movement sequence as an integrated unit. The completion of one finger controller automatically triggers the next finger controller. **(b)** A hierarchical controller represents the movement sequence across multiple layers that interact to produce the sequence of movements. The finger controllers represent the specific muscle commands for each of the fingers and are responsible for finger press execution. The sequence controller commands the finger controllers to initiate movements. In this particular model, the finger controllers provide internal feedback to the sequence controller when the finger press is completed. However, the next press may be initiated at a different time from the occurrence of the internal feedback.

While both models would predict a modulation of the press that is perturbed, they differ in how subsequent presses would be affected. In the single motor program model, an acceleration or delay of a single movement element will shift the subsequent presses accordingly. In contrast, in the hierarchical model, the influence of a local sensory perturbation on a single finger could differ from the influence on subsequent presses. How exactly subsequent presses are influenced depends on how feedback is communicated from the lower-level finger controllers to the higher-level sequence controller (Kiebel et al. 2009), and how the sequence controller uses the feedback. By comparing the influence of a sensory feedback perturbation across finger movements of a sequence, we are able to gain novel insights into how sensory feedback is used in this organization.

## Methods

### Participants

Twenty-six participants were recruited for this study (11 males; ages 18 to 44; mean age 25.5 [± 7.25]). All participants were right-handed (self-declared) and completed informed consent. On average participants had received 6.44 (± 7.25) years of musical training based on their longest played instrument, with 57% having at least one year of piano playing experience. The study protocol was approved by the ethics board of the University of Western Ontario and all participants gave their signed consent before starting the study.

### Apparatus

To test participants, we used a custom-built five-finger keyboard (Fig. 2a). The keys were not depressible but a force transducer (FSG-15N1A, Sensing and Control, Honeywell) was mounted underneath each key measuring isometric force production with a repeatability of <0.02 N and a dynamic range of 16 N (Wiestler et al. 2014; Wiestler and Diedrichsen 2013; Yokoi et al. 2017). The digital sampling rate of the measured force was 200 Hz. Additionally, each key was equipped with a linear resonant actuator (LRA, LVM061930B-L20, Jinlong Machinery & electronics Inc.) that provided haptic feedback during the experiment. LRAs vibrate at a frequency between 200 and 250 Hz. In our application, a haptic controller creates a specific waveform to elicit the click sensation. The haptic stimulation was produced by a haptic motor controller (DRV2605L, Adafruit Industries LLC) that produces a computer-controlled click/vibratory sensation that feels similar to the sensation experienced from smartphone keys or trackpads on laptops (see the DRV2605L dataset for more information regarding the specific waveform).

**Figure 2.**
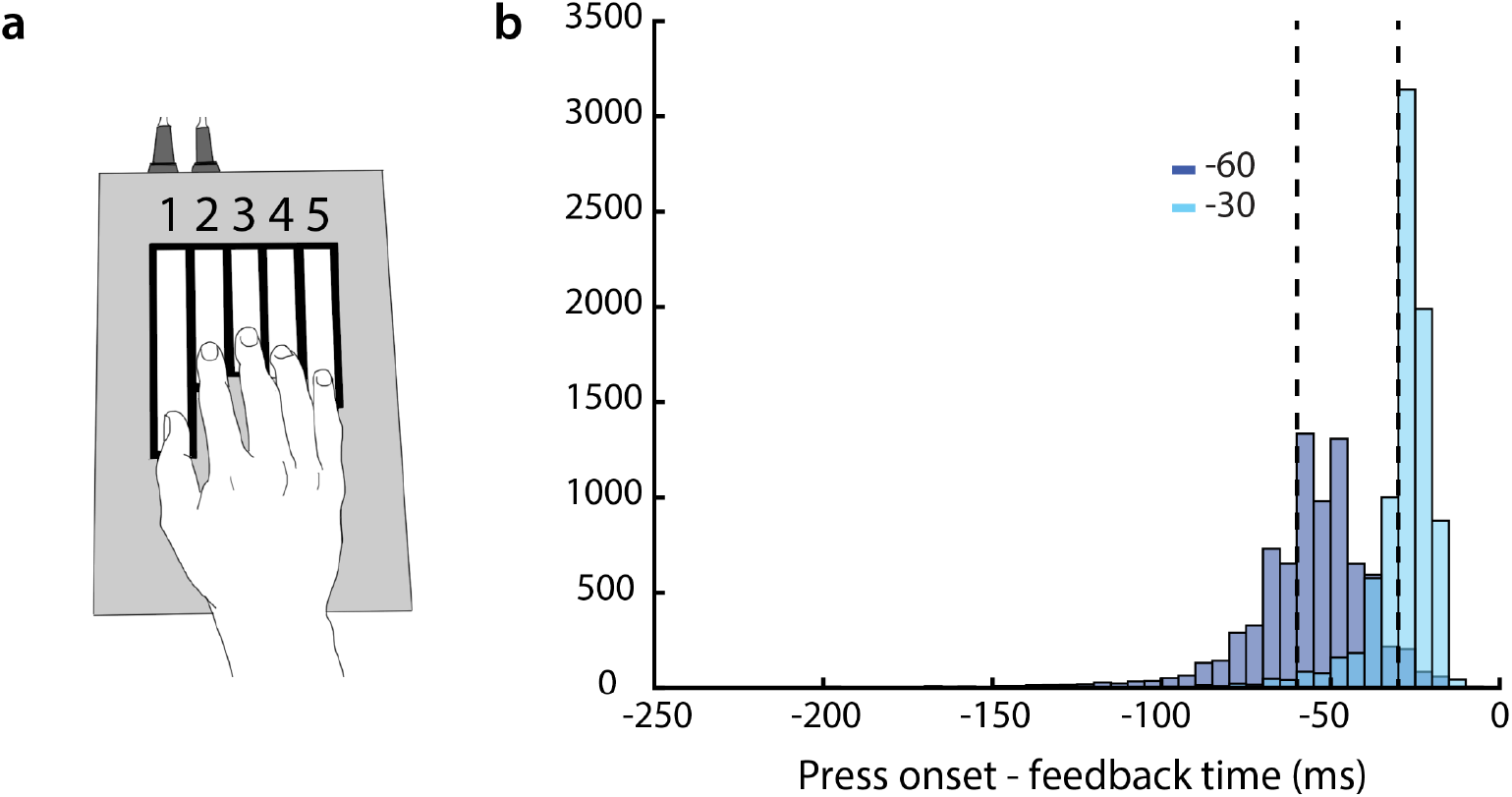
Apparatus and achieved time advancements of feedback. **(a)** Isometric keyboard-like device. Each key was associated with a number (these numbers were not shown to the participants but verbally explained). **(b)** Histogram of the time intervals between feedback presentation and the actual press onset for the two advancement conditions. Vertical doted lines indicate -30 ms and -60 ms.

### Discrete sequence production task

Participants performed a discrete sequence production task (DSP), executing sequences of 11 keypresses as fast and as accurately as possible. Participants were instructed to move as fast as possible while maintaining an error rate of under 15% for each block of trials. Each finger was associated with a number (thumb = 1, index = 2, middle = 3, ring = 4 & little = 5). Each trial began with the presentation of a sequence of numbers on a computer screen (white font). A trial was deemed completed after 11 finger presses were executed. The numbers stayed on the screen throughout execution.

Participants performed three sequences in total that were randomly presented to the participant. None of the sequences had directly repeating numbers (i.e., 33 or 44). The same three sequences were used for all participants; however, the presentation order was randomized across participants. Each block consisted of 39 trials and each sequence was presented 13 times during a block.

The force magnitude applied to each key by the participant was displayed as five lines on an LCD monitor, where each line height indicated the amount of force applied to the corresponding key. When the force on a key exceeded 1.5 N, the keypress was registered and the feedback was triggered. Some co-articulation between fingers emerged as the next key could be pressed before the previous key was released.

When participants pressed the correct key, the visual cue on the screen turned green, a short pleasant auditory sound could be heard (each key was assigned a specific tone that was different from the rest) and a small click could be felt on the finger. If, however, an incorrect key was pressed, the visual cue changed to red, a lower-pitch sound could be heard (same across keys), and a click (same for accurate and incorrect press) could be felt.

For each completed trial participants received points based on their performance. If the participant pressed all keys correctly and their median movement speed (MT – the time between the first press and last release) was within 95% to 110% of the current speed threshold (MT threshold) they received one point. If they correctly executed the sequence and their median movement speed was faster than 95% of the current MT threshold they received three points. If they pressed one or multiple keys incorrectly or their median speed was slower than 110% of their MT threshold they received zero points. At the end of a block, we provided participants with feedback regarding their error rate, median speed (MT), points obtained for the current block, and total points obtained across the session. To motivate participants to improve their performance throughout the sessions, we first set the MT threshold at 10 s at the beginning of each session and then adjusted it by lowering it to the median MT of a given block if the participant had a lower median MT compared to the current MT threshold and if their error rate was below 15%.

### Feedback manipulation

The first three blocks in each session were completely unperturbed, meaning no feedback perturbation was presented. In each of the following blocks, we perturbed 24 trials out of the 39 trials. Participants completed a total of 74 blocks over the four days of training. On these perturbation trials, we either advanced or delayed the haptic, visual, and auditory feedback by 30 or 60 ms on one of the 11 key presses. To generalize our findings across fingers and press location within the sequence, we chose two fixed positions within each sequence where feedback perturbations were given. This also reduced the potential predictability of the perturbation location in each sequence. In sequence 1, we gave the feedback perturbation either at position 6 (finger 5) or 9 (finger 4), in sequence 2 at positions 4 (finger 2) or 7 (finger 1), and in sequence 3 at positions 5 (finger 4) or 8 (finger 3). In total, we presented the perturbation at six different sequence positions across all sequences.

For the advanced feedback conditions, we used an algorithm to predict when the feedback had to be given to occur either 30 or 60 ms before press onset (the time at which the force on the key exceeded 1.5 N). This prediction was updated in real-time every 2 ms during trial execution. This prediction was based on three factors: the current force, the current force change (numerical derivative based on three time points), and the time since the last press onset. We separately trained this predictive model for each subject, sequence position, and delay condition (−30 ms or -60 ms) using a logistic regression. This was done twice in each session. The first time we fit the model on the data from the first three blocks, using the unperturbed trials as training data. To account for speed changes during the session, we repeated the estimation in the middle of the session based on the unperturbed trials of all previous blocks (excluding the three initial blocks and at least six blocks of trials). The predicted outcome variable was zero if it was too early to present feedback and one if it was too late. Feedback was provided once the predicted probability exceeded 0.5.

This approach led to an average time advancement of 29.3 ms (SD: 11.4 ms) for the -30 ms advancement condition and an average of 57.9 ms (SD: 23.3 ms) advancement for the -60 ms condition (see Fig. 2b).

On the advanced trials, participants could press the next key as soon as the feedback was presented on the current press, meaning they were allowed to press the next key before reaching the press threshold for the perturbed press. This led to an average of 2.36% (SD: 1.55%) of the advanced trials not reaching the press threshold. We excluded these trials from our analyses. Our analyses centred on calculating time intervals between specific press landmarks and the press onset of the perturbed press. In these trials the press onset was absent and thus we were unable to perform the same analyses.

In the delay conditions feedback was withheld upon reaching the press threshold, and instead presented 30 or 60 ms after press onset. However, in the delay conditions participants were not required to wait for the feedback to be presented before moving on to press the next press. This was important as participants did not have to take the feedback perturbation into account and could potentially perform the sequences just as fast as when no perturbation was present.

### Experimental Procedure

Participants completed four sessions that lasted approximately 1.5 hours each depending on how fast the participant was able to complete the required blocks of trials. Participants completed one session per day and the four sessions were scheduled over a time span of approximately two weeks. We encouraged participants to take breaks between blocks as necessary and offered a longer break in the middle of the experimental session. The participants were told that the goal was to perform the sequences as accurately and fast as possible. At the end of the four sessions, we asked participants several questions about their experience that became more and more specific (see Appendix 1). This questionnaire was used to determine whether participants were conscious of the experimental manipulation. Only two participants expressed clear conscious knowledge of the experimental manipulation, while the rest of the participants did not notice the manipulation. The performance of these two participants was similar to the performance of the other participants and therefore were not excluded from the analyses. Overall, the majority of participants were not consciously aware of our experimental manipulation, and hence we believe that they did not change their behaviour consciously.

### Statistical Analysis

For each trial, we calculated the overall movement speed (movement time/MT) between the onset of the first press (first time it reached the press threshold) and the release of the last press (force fell below 1 N). Additionally, we found five landmarks (Fig. 3a) for each press: early onset (*EO* - when force first was great or equal to 0.75N), onset (*O* - when force first was equal or exceeded 1.5 N), peak (*P* - time at highest force – between onset and late release), release (*R* - when the force first fell under 1.5 N after peak), and late release (*LR* - when force first fell under 0.75 N after onset). All analyses were done relative to the onset of the perturbed press (or for unperturbed trials, the matching unperturbed press in the same sequence). We analyzed the relative timing of the landmarks on the perturbed press (+0), and the two presses after the perturbed press (+1 & +2).

**Figure 3.**
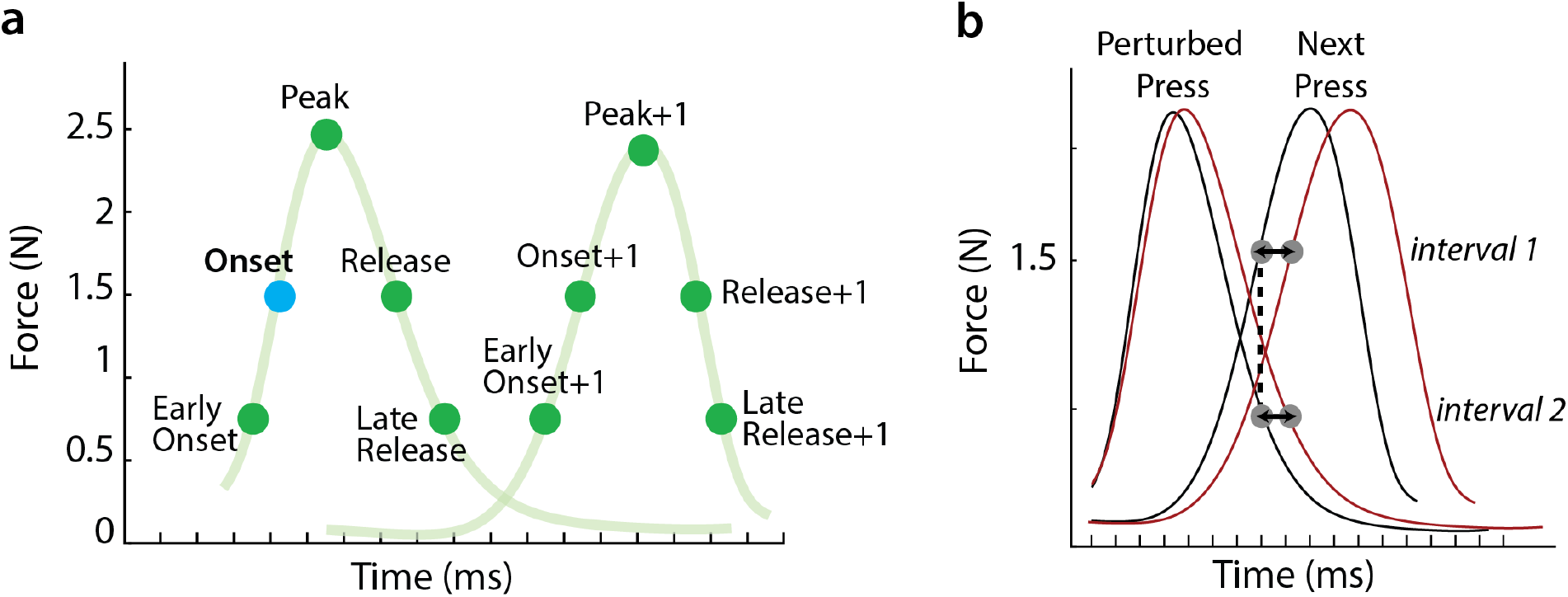
Calculation of feedback differences across presses and landmarks. **(a)** For our analyses we calculated time intervals between the onset of the perturbed press (blue onset dot in the figure) and different force landmarks (green dots) on the perturbed press as well as on subsequent presses (indicated with +1). We chose five specific force landmarks on each press: Early Onset (>=0.75 N), Onset (>=1.5 N), Peak (maximum N between onset and release of press), Release (first time <1.5 N after onset), and Late Release (first time <0.75 after onset). **(b)** We choose a single time point (onset of next press) and compared how the perturbation affected this time point across presses. The black line indicates unperturbed trials and red lines represent perturbed trials.

All analyses were performed using custom-written code in MATLAB (The MathWorks) and the dataframe toolbox (github.com/jdiedrichsen/dataframe). We excluded any error trials from our analyses, as well as trials in which the press was delayed by more than 100 ms after the advanced feedback was given, as we believe that this could either suggest conscious awareness or an incorrect estimation from our algorithm that predicts when feedback should be given. We analyzed the data using paired one and two-sample t-tests that were based on clear a priori predictions and we chose a probability threshold of p<0.05 for the rejection of the null hypothesis.

To estimate how quickly participants reacted to the delayed feedback by adjusting the perturbed press, we conducted a change point analysis. We first calculated the difference between the average force curves for the delayed trials (+30ms or +60ms) and unperturbed trials from 20 ms before press onset and 240 ms after onset. Using the data before the occurrence of the peak difference between the two curves, we estimated the time point when the difference started to emerge. We modelled the difference as a piece-wise linear function with a change point of b_0_ between the two segments.

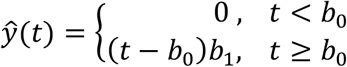

where ŷ (*t*) is the predicted force values for time *t, b*_0_ is the chosen change point and *b*_1_ is the slope of the function. Using the function fminsearch in MATLAB, we found the values for *b*_0_ and *b*_1_ that minimized the sum of squares of the difference between observed and fitted data.

The single motor program hypothesis predicts that the perturbed press (+0) and the press following the perturbed press (+1) would be delayed or accelerated (relative to an average unperturbed press) by the same amount (Fig. 3b). To test this idea, we examined the difference in the effect of the perturbation at a singular point in time across the consecutive presses (i.e. a point in time where the force curves of the presses overlap). We first chose a landmark at a time when the force curves of the two presses overlapped. At the end of training, this overlap was clearly observed at the onset (1.5 N) of the +1 press for the unperturbed trials, which we chose as our reference landmark. On unperturbed trials, we then found the average force for the +0 press, which defined our matching landmark (i.e. that occurred at the same point in time; see Fig. 3b). We then calculated the effect of the perturbation on these two landmarks. The single motor program hypothesis predicts that both landmarks would be delayed by the same amount of time (relative to an unperturbed press). In contrast, a difference in delay (positive or negative) between the +1 press and the +0 press would indicate that the effect of the perturbed feedback was not the same for the two presses.

### Control experiment

In a separate experiment, we probed to what degree the modality of the sensory feedback (auditory, haptic and visual) had differential effects on participants’ performance. We recruited 48 participants for this experiment. They were assigned to one of the three feedback groups (auditory, haptic or visual) at the beginning of training based on an algorithm that matched participants’ speed, calculated as the time between the onset of the first press to the release of the last press (MT). This was done to ensure that the groups had similar average speeds at the start of the experiment. Participants only received one type of feedback throughout the study (how each feedback was given was the same as described in the experimental design above). When an incorrect finger press occurred, all groups saw the visual cue on the screen turn red to make it easier for them to know where they made the error in the sequence. Participants practiced four different sequences (three were the same as in the main experiment) for five days on the same keyboard-like device. Press threshold was 1 N. Because of the difference in press threshold we adjusted our landmark criteria for this experiment: early onset (*EO* - when force first was great or equal to 0.6 N), onset (*O* - when force first was equal or exceeded 1 N), peak (*P* - time at highest force – between onset and late release), release (*R* - when the force first fell under 1 N after peak), and late release (*LR* - when force first fell under 0.6 N after onset). Feedback perturbations were given on a single press within the sequence at two possible locations (similar to the main experiment but the locations were not identical). In this experiment, we only perturbed participants’ feedback by delaying it by 80 ms. The rest of the experimental design was identical to the main experiment (point system, threshold change etc.). As in the main experiment, most participants were unaware of the perturbation when asked about it using a questionnaire at the end of the sessions.

## Results

### Feedback perturbations cause directionally specific behavioural adjustments to the perturbed finger press

To investigate how sensory feedback is used during the execution of fast finger sequences, we used transient perturbations of the sensory feedback that indicated the successful pressing of a key. The perturbation was only applied to a single press within a sequence. Participants practiced three different sequences over four days. If sensory feedback is used to control the near-isometric keypress, the delay and advancements of feedback should prolong or shorten the ongoing press, respectively.

The group average force traces (Fig. 4a) indicated that even though each finger press was completed within ∼300 ms, participants indeed reacted to the feedback perturbation by extending or shortening the ongoing press. To quantify this effect, we calculated the time interval between the onset (first time >=1.5 N is reached) and the peak (onset-peak) of the perturbed press (Fig. 4b onset-peak), as well as the interval between the onset and the release (first time <1.5 N after onset; Fig. 4b onset-release). On day 1, both the +30 ms (t_(25)_ = 11.189, p= 1.59e-11) and the +60 ms delay condition (t_(25)_ =4.969, p= 2.02e-05) resulted in a longer onset-peak intervals. Similar effects can also be seen on the interval between onset and release (+*30 ms*: t_(25)_ = 6.630, p = 3.01e-07, +*60 ms*: t_(25)_ = 5.963, p = 1.58e-06). For the time advanced feedback conditions, the onset-release intervals on day 1 were shortened in response to perturbations (onset-release *-30 ms*: t_(25)_ = 5.308, p = 8.42e-06; *-60 ms*: t_(25)_ = 4.291, t= 3.78e-10). These results suggest participants used sensory feedback indicating the successful pressing of a key to finely control the duration of the force production.

**Figure 4.**
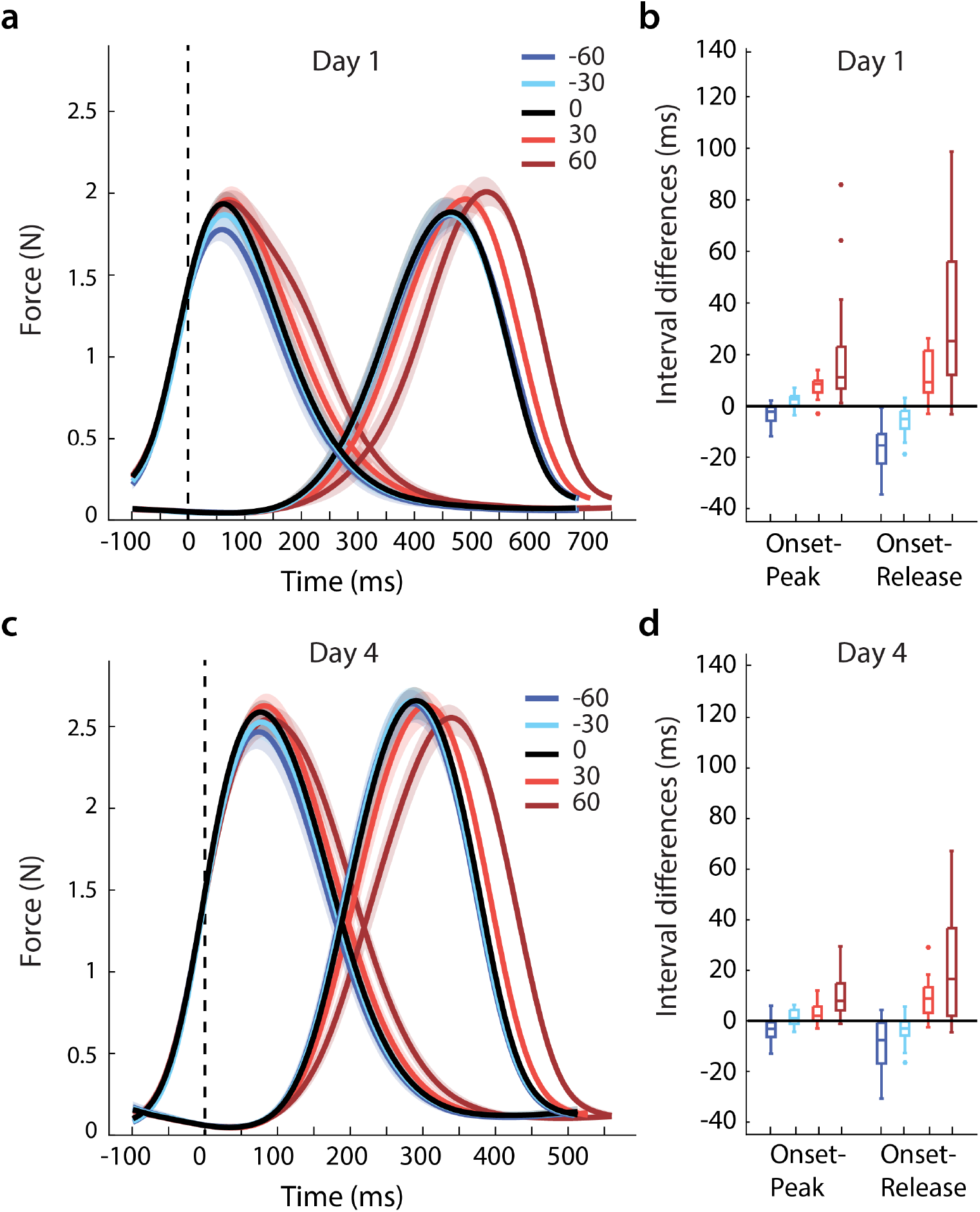
Effects of perturbation on perturbed press and subsequent press. **(a)** & **(b)** Average force traces for day 1 and 4 and the following press interpolated and standardized to the average time of each condition. Dotted line indicates press onset, for which the sensory feedback was shifted in time. Error bars represent the standard error of the mean across subjects. **(c)** & **(d)** Differences between the onset-to-peak and onset-to-release intervals of perturbed and unperturbed trials for day 1 and day 4.

### Perturbation effects diminish but do not disappear with training

Does feedback control still play a role in movement execution the end of training? If the motor system uses sensory feedback to control the execution of extensively practiced finger movements, we expect the feedback perturbation to still impact the duration of the press at the end of training. Indeed, this was what we found (Fig. 4a vs. 4c). Specifically, both delay conditions showed larger onset-peak intervals (*+30 ms*: t_(25)_ = 5.963, p = 1.17e-04; *+60 ms*: t_(25)_ =6.420, p= 5.05e-07) and onset-release intervals (*+30 ms*: t_(25)_ = 6.143, p = 1.01e-06, *+60* ms: t_(25)_ = 5.082, p = 1.51e-05) compared to the unperturbed condition on day 4 of training (Fig. 4d). Similarly, shorter onset-release intervals were observed for the advancement conditions (day 4 onset-release *-30 ms*: t_(25)_ = 3.774, p = 4.46e-04, *-60 ms*: t_(25)_ = 4.785, p= 3.26e-05). The finding of a clear adjustment of the perturbed press at the end training suggests that even skilled performance is controlled by sensory feedback.

While the overall effect was clearly present across all days, the effect caused by the large perturbations reduced by ∼40%. Specifically, the difference between perturbed and unperturbed onset-release interval reduced from day 1 to day 4 for the +60 ms (−38%, t_(25)_ = 2.502, p = 0.019) and the -60 ms condition (−40%; t_(25)_ = -3.859, p = 7.106e-04). While the overall effect also reduced for the smaller perturbations, these changes were not significant (+30 ms: -29%, t_(25)_ = 1.848, p = 0.076; -30 ms: -35%, t_(25)_ = -1.639, p = 0.113). This suggests that some transition from feedback to feed-forward control took place in our task with practice.

### Perturbations lead to reactions within 80ms

How quickly is sensory feedback taken into account to control the ongoing finger press? To estimate this, we first calculated a difference curve between the average force traces of the delayed perturbation conditions and the unperturbed condition for each participant. We then used a change point analysis (see methods for details) to estimate the time at which the difference curve was impacted by the feedback delay. On day 1 in the +60 ms delay condition, it took an average of 106.4 ms (95% CI [97.77, 115.03]) after press onset for participants to show a divergence between the two force traces. For the +30 ms delay condition, the difference started at 77.3 ms (95% CI [64.65, 90.04]). For day 4, our estimate of adjustment onset for the +60 ms condition was 92.5 ms (95% CI [83.04, 101.97]), faster than day 1 (t_(25)_ = 2.085, p = 0.047). The estimate for the +30 ms condition was comparable to day 1(mean: 67.5; 95% CI [46.32, 88.72]; t_(25)_ = 0.738, p = 0.467). Overall, the adjustment of the ongoing press to the delayed feedback was consistently very fast.

### Subsequent presses are delayed irrespective of perturbation direction

So far, we have established that sensory feedback about the keypress is used to control the finger that produces the press, even during fast performance after extended training. Next, we investigated how the subsequent presses are impacted by the perturbation. This provides us with an opportunity to compare different models of how skilled movement sequences are organized.

To visualize how the perturbations influence both the current and subsequent presses, we plotted the timing of five events (early onset, onset, peak, release, late release, see Methods) for the perturbed and the two subsequent presses across the four sessions (Fig. 5). As the independent variable (i.e. x-axis) we plotted the group-averaged time estimates of these landmarks for the non-perturbed trials relative to the onset of the perturbed press (0 ms). As the dependent variable (i.e. y-axis) we plotted the change in the average time interval relative to the unperturbed condition. Each press is indicated by a line that connects the five corresponding landmarks.

**Figure 5.**
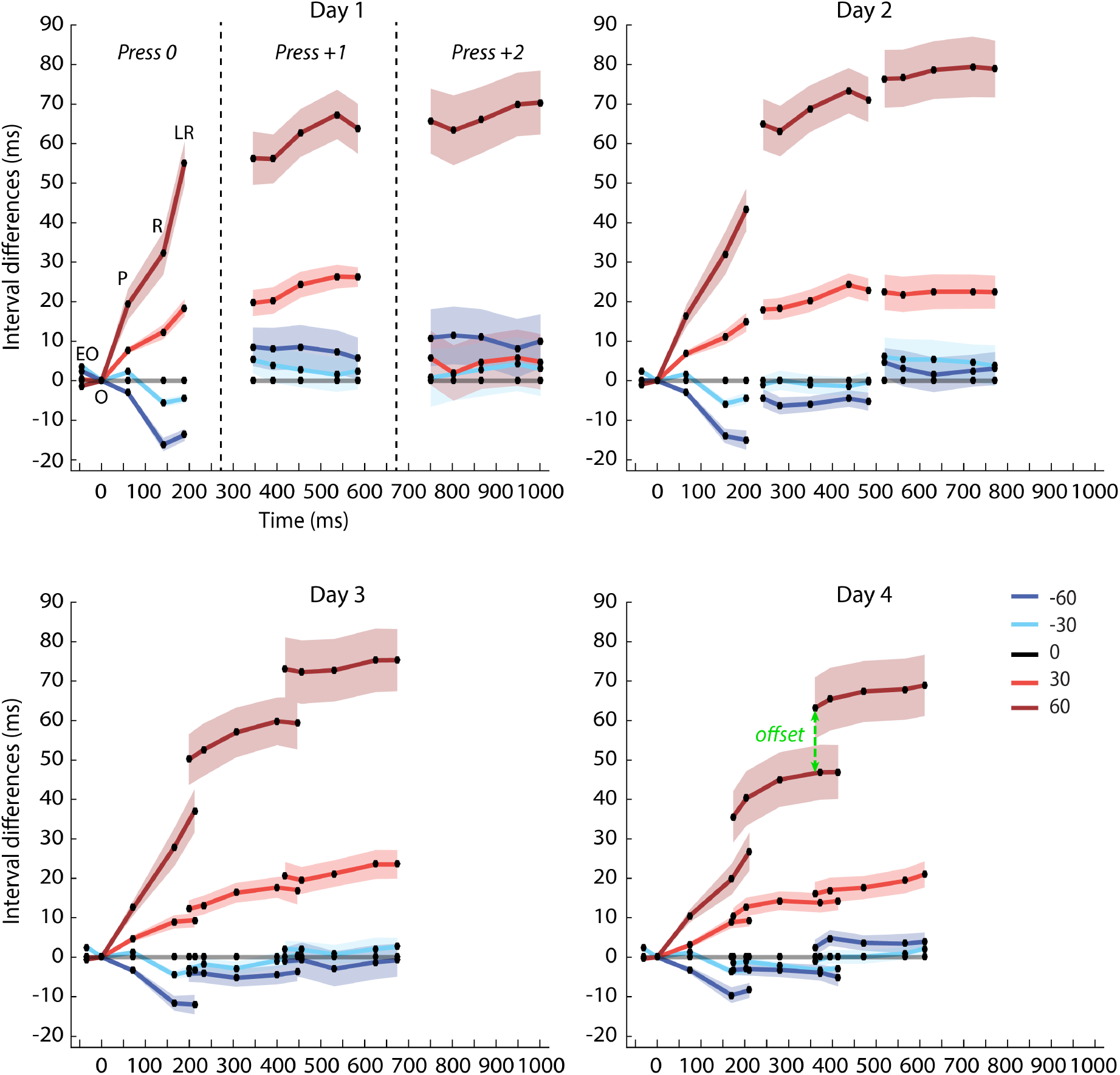
Effects of feedback perturbation on the perturbed press (press 0) and subsequent finger presses (Press +1, +2) across feedback conditions and training days. Five landmarks (EO: early onset, O: onset, P: peak, R: release, LR: later release) are plotted per press (see methods). The x-axis shows the average time of occurrence of the landmark on unperturbed trials relative to the onset of the first press. The y-axis shows the time interval differences between the perturbation conditions and the unperturbed condition on the particular landmarks. Landmarks belonging to a finger press are connected by a line. Anything above the 0 line indicates that the perturbation resulted in longer time intervals (i.e. slower) compared to when no perturbation was present, whereas everything below the line indicates shorter time intervals (i.e. speed-up). The different panels indicate the different training sessions (i.e. days). Day 4 shows how we tested the offset between presses, with an example of the 2^nd^ to 3^rd^ press for the +60ms condition. Error bars represent the standard error of the mean across participants.

The feedback perturbations impacted not only the execution of the current press, but also of subsequent presses. On the first day of training, both the +30 ms perturbation (t_(25)_ = 6.055, p= 2.51e-06) and the +60 ms perturbation (t_(25)_ =9.078, p= 2.177e-09) delayed the onset (interval onset-onset+1) of the next press relative to when no perturbation was present (i.e. red lines vs. grey line at zero). Moreover, the delay of feedback impacted even the onset of the press two positions after the perturbation (+*60 ms*: t_(25)_ = 7.172, p= 8.11e-08). In contrast, time advancements did not alter the timing of subsequent presses relative to the unperturbed trials (onset-onset+1: *-30 ms*: t_(25)_ = - 0.904; p = 0.375; *-60 ms*: t_(25)_ = -1.488, p = 0.149). This pattern of results provides new insights into how feedback is used in the control and representation of skilled movement sequences (as outlined in the introduction, Fig. 1).

If trained sequences are encoded as a single motor program (Fig. 1a), the control of one finger directly influences the control of the subsequent finger. This prediction becomes directly testable when there is considerable overlap, i.e. coarticulation, across different finger presses. Such coarticulation was observed on days 3 and 4 (Fig. 5; where the onset of the second press roughly occurred at the same time as the release of the perturbed press). For such overlapping presses, the single motor program hypothesis would predict that the relationship between the release of the perturbed press and the onset of the next press in the sequence will be the same, even if the entire motor program is sped up or slowed down. In other words, the effect of the perturbation should be the same for simultaneous events on two overlapping presses. To test this idea, we used the data from the last day of training. We compared the effect of the perturbation on the onset of the next press (onest+1 Fig 5) with its effect on the perturbed press at the same point in time (see Methods for detail). We found a significantly longer delay for the subsequent press in comparison to the perturbed press for the +60 condition (t_(25)_ = 2.522, p = 0.018). This effect can be seen as an offset between the end of the line for the perturbed press and the onset of the line of the subsequent press in Figure 5 (day 4). A similar offset between presses was also present between the second and third press after the perturbation (t_(25)_ = 3.429, p = 0.002). These additional delays across presses resulted in an overall slower execution speed for the entire sequence (MT; day 4: *+60 ms*: t_(25)_ = 5.828, p = 4.456e-06). These findings provide clear evidence against the idea that the sequence is represented as a single motor program after training. Rather it argues for a hierarchical organization (Fig. 1b), in which the effects on the subsequent finger presses can differ from the effect on the perturbed finger.

The participants’ reactions to the other perturbation conditions provide us with more detailed insight into how feedback is considered in this hierarchical organization. Similarly to what we have observed for the +60 ms delay condition, an offset between the different presses was also observed for time-advancement of the feedback by -60 ms (dark blue in Fig. 5), although this effect did not reach significance (t_(25)_ = 2.043, p = 0.052). Nevertheless, the offset was significant when comparing the second and third press after the perturbation (t_(25)_ = 3.877, p = 6.799e-04). In the -60 ms perturbation condition, these additional offsets did not result in a significant slowdown of the overall sequence speed (Day 4: t_(25)_ = -0.858, p = 0.399), suggesting that the additional delays of subsequent presses were cancelled out by the speed-up on the perturbed press. In contrast to the ±60 ms feedback perturbations, no clear offset was present for the ±30ms perturbation condition (Fig. 5 – light blue and light red). Indeed, the comparison did not reach statistical significance for either time delay (+*30 ms*: t_(25)_ = 0.882, p = 0.193) or advancement perturbation (*-30 ms:* t_(25)_ = 0.589, p = 0.281). In sum, for larger but not for smaller perturbations participants delayed subsequent presses after the occurrence of a perturbation, irrespective of whether the sensory feedback was advanced or delayed.

Overall, our findings suggest a hierarchical organization in which sensory feedback acts in two qualitatively different ways. First, the timing of the feedback *directionally* either lengthens or shortens the perturbed key press. Second, the occurrence of a perturbation also appears to act in a *directionally non-specific* manner slowing down the execution of future presses. This effect was stronger for larger (60) compared to smaller (30) perturbations but did not depend on the direction of the temporal shift.

### Rapid behavioural adjustments are caused by haptic feedback

Finally, we investigated to what degree the effects observed in the main experiment were due to the perturbation of haptic, visual, or auditory feedback. To test this, we conducted a control experiment, in which a separate set of participants was assigned to one of three experimental groups, with each group receiving only one of the three types of feedback (auditory, visual or haptic). As in the main experiment, we delayed the feedback on selected finger presses within the sequence. In this case, we only chose a single perturbation condition (delay +80 ms) and participants practiced the task for five days. Examining the effect of the delay on the perturbed press (see Fig. 6), we found that only the haptic group demonstrated a significantly longer onset-peak interval following the perturbation both in the beginning (Day 1: t_(15)_ = 2.980, p = 0.009) and towards the end of training (Day 4: t_(15)_ = 3.579, p = 0.003). Neither the visual (Day 4: t_(15)_ = 0.901, p = 0.382) nor the auditory group (Day 4: t_(15)_ = 1.060, p = 0.306) showed a significant effect of the feedback perturbation on the onset-peak interval. These results clearly show that the rapid adjustments of the ongoing press were driven by haptic feedback from the fingertip.

**Figure 6.**
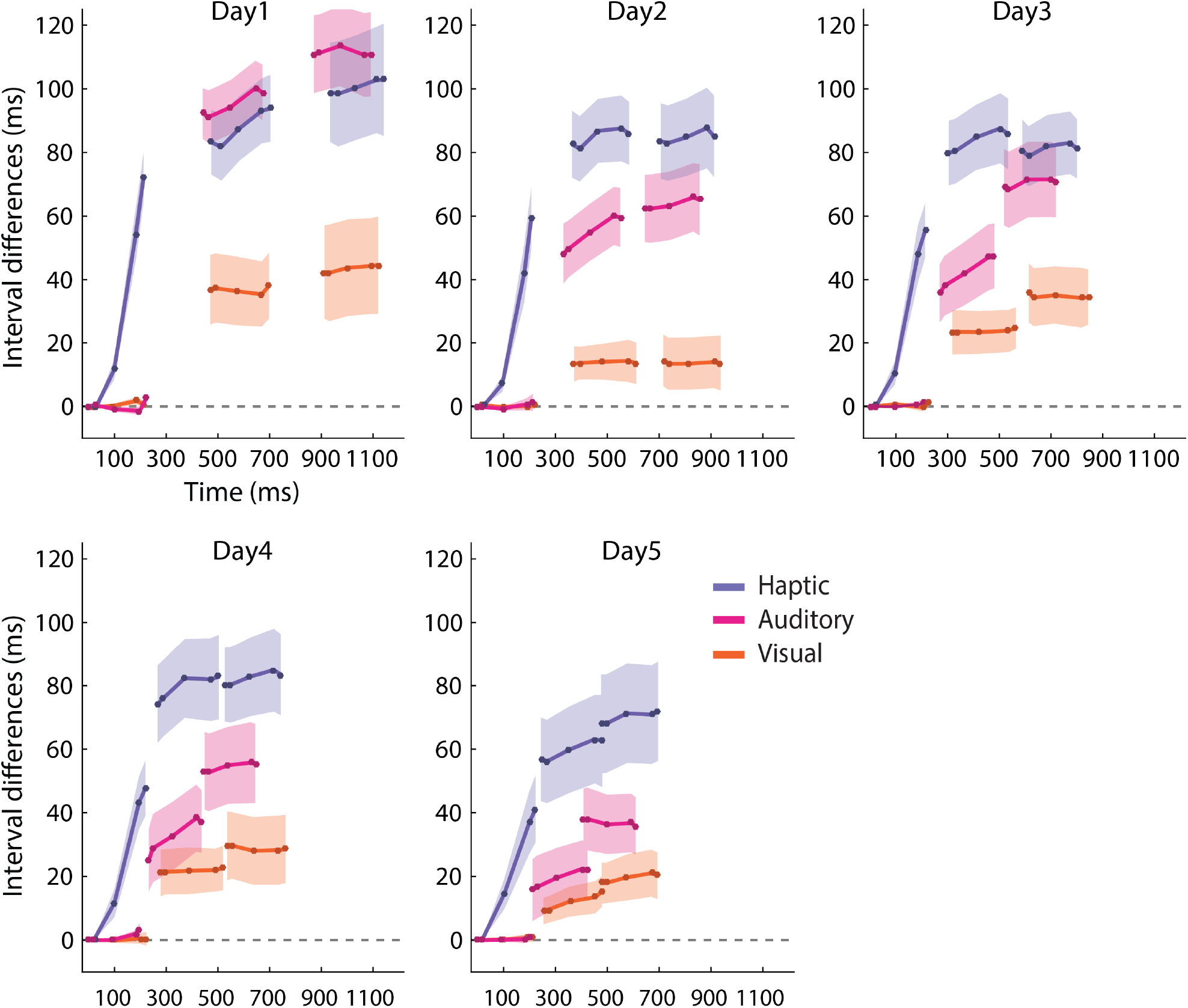
Effect of feedback perturbation for haptic, visual and auditory groups in control experiment across training days. As in Figure 5, five landmarks per press (connected by a line) are plotted. The control experiment only had +80ms perturbations, but each group received only one type of feedback. The different panels indicate the different training sessions (i.e. days). The error bars represent the standard error of the mean across participants for each group.

### Delay of subsequent presses arises from all three feedback modalities

In contrast, the delay of subsequent presses was observed for all three feedback modality groups. Consistent with the effect on the perturbed press, the delay of the onset of the press following the perturbation (+1, averaged across days 2-5) was largest in the haptic group (69 ms, t_(15)_ = 6.890, p = 5.146e-06). However, both the auditory group (35 ms, t_(15)_ = 4.888, p = 1.971e-04), as well as the visual group (19 ms, t_(15)_ = 4.828, p = 2.214e-04), showed a clear delay in the onset of the subsequent press, even though no such effect was observed on the perturbed press (Fig. 6). This result suggests that the delay we observed on the subsequent presses in our main experiment could be induced by the perturbations in each of the three feedback modalities.

## Discussion

In this study, we used small transient feedback perturbations to probe how sensory feedback is used in the control of fast finger movement sequences. Specifically, we examined how sensory feedback modulates the execution of skilled finger movements across four days of training, and how feedback differentially affects the execution of the ongoing press and subsequent presses.

### Sensory feedback rapidly modulates movement execution of the perturbed press

Throughout training, we found clear evidence of rapid behavioural adjustments of the finger press that received the perturbation. This result illustrates the continuous integration of sensory feedback when controlling skilled finger movements. Participants adjusted their ongoing behaviour even though our task was designed so that it could be accomplished without considering the feedback. The keypresses were isometric and participants simply needed to exceed a specific force threshold. Nonetheless, participants adjusted their behaviour based on the perturbation.

Furthermore, we found that the effects of the perturbation were directionally specific: The delay in sensory feedback resulted in a lengthening of the perturbed press, whereas a time advancement resulted in a shortening. Previous studies have primarily investigated feedback delays (Furuya and Soechting 2010; Howell and Archer 1984; Sakata and Brainard 2006; van der Steen et al. 2014) but have rarely advanced participants’ feedback (Repp 2002; Wing 1977). By including both feedback delays and advancements we provided evidence of the directional nature of sensory feedback integration in fast finger movements.

The reaction to the delay of haptic feedback was very fast and occurred within 60-90 ms after the expected time of the feedback. This finding is consistent with previous reports that demonstrate responses between 65-110 ms following a haptic input (Abbs et al. 1984; Pruszynski et al. 2016; Scott 2016). In contrast, auditory and visual feedback alone did not elicit a strong reaction on the press, consistent with the fact that the fasted reactions to changes in these two modalities are noticeably slower (Burnett et al. 1998; Day and Lyon 2000; Howell 2004; MacKenzie and Marteniuk 1985; Smith and Bowen 1980; Veerman et al. 2008). Therefore, by including a haptic feedback condition we were able to show the very rapid integration of sensory feedback in the execution of a finger press.

### Shift from feedback to feed-forward control with learning

While the feedback perturbation still impacted the execution of the perturbed press on the last day of practice, we did find that the effect reduced by approximately 40% with training. This observation is in line with previous research that observed a shift from feedback to feed-forward control with training (Pew 1966; Seidler-Dobrin and Stelmach 1998). It has been suggested that feedback plays an important role in the initial phases of acquiring a novel motor skill, but its importance decreases, and potentially even disappears altogether, with prolonged training (Pew 1966; Pratt et al. 1994; Schmidt 1975; Schmidt and McCabe 1976; Seidler-Dobrin and Stelmach 1998). The main theoretical idea here is that, as we acquire an accurate internal representation of the instructed movements, sensory feedback becomes less necessary for execution (MacNeilage and MacNeilage 1973; Schmidt 1975; Seidler-Dobrin and Stelmach 1998). Alternatively, participants potentially learned that the large deviations of the sensory feedback were irrelevant for overall performance and therefore could be ignored (Wei and Körding 2009).

### Distinct feedback processes govern the control of rapid finger movement sequences

Our second goal was to understand how sensory feedback is being taken into account in the control of a complex motor sequence. Models of sequence representation fall between two opposing extremes: A single, integrated motor program, and a hierarchical organization (Fig. 1). By examining how feedback is integrated across multiple finger presses, we were able to investigate the underlying organizational structure and how feedback is integrated across the different layers.

We found that the feedback perturbation on a single press also affected the execution of subsequent presses, both at the beginning and at the end of training. Important, the reaction to the feedback perturbation was different for the perturbed and subsequent presses. This finding argues against the idea that after prolonged training a movement sequence is represented as a single motor program (Keele 1968; Rozanov et al. 2010), in which each finger is affected in the same way by the perturbation. Instead, our results more closely align with the idea of a hierarchical organization (Rosenbaum et al. 1983), in which the sequence is controlled through the interaction of different layers that control sequence execution. One possible organization is a two-tiered structure (Fig. 1b), in which a sequence controller is positioned at the highest level representing the specific order of movements and commanding the next layer of finger controllers, which in turn are responsible for the control of specific finger movements. Our results suggest three characteristics of this proposed two-tiered control structure:

First, sensory feedback from the finger itself is continuously relayed to the finger controller which then impacts the ongoing movement execution in a directional specific manner. Second, upon receiving the sensory feedback signifying press completion, the finger controller issues a completion signal to the sequence controller. Our finding that feedback not only impacts the ongoing press but also subsequent presses, suggests that information is relayed across all hierarchical levels. Third, we found that both feedback advancements and delays led to an overall slower initiation of the next finger in the sequence. This slowing possibly reflects a cautionary measure: The sequence controller may compare a prediction of when a completion signal is expected versus when it is received and, upon detecting a mismatch, delays the execution of the next press to ensure the successful completion of the sequence. We also found that only the two larger sensory feedback perturbations led to a significant delay, suggesting that the cautionary response is proportional to the amount mismatch between expected and received feedback from the lower-level controller. Additionally, the sequence controller also showed a reaction to a delay or time advancement in auditory and visual feedback, which did not influence the local press, indicating that the sequence controller also has direct access to sensory feedback signalling whether the goal of an action has been achieved.

Previous research studying time delays and advancements of an external pacing signal in a synchronization paradigm (Furuya and Soechting 2010; Repp 2000; Wing 1977) also have shown evidence for feedback adjustments in a hierarchical sequence controller. In contrast to our experiment, in which a feedback perturbation led to a delay irrespective of direction of the perturbation, these adjustments were targeted at bringing the finger presses back into synchronization with the metronome (Furuya and Soechting 2010; Repp 2000). In our paradigm, the timing was not constraint by a metronome, such that the task goal was not to preserve a rhythm. The difference in findings between these two paradigms make it likely that the reaction of the sequence controller to feedback perturbation will strongly depend on the task goal.

### Conclusion

In this study, we demonstrated that sensory feedback is continuously used to adjust movement execution, but that the extent of this integration diminishes with training. Haptic feedback drove the effects we observed on the perturbed press, whereas the effects across the remaining movements in the sequence were impacted by the perturbation in all three feedback modalities. Lastly, we demonstrated distinct types of feedback processes involved in the hierarchical control of skilled finger sequences.

## Acknowledgments

This work is supported by a Canada First Research Excellence Fund (BrainsCAN) to Western University, a Natural Sciences and Engineering Council of Canada (NSERC) Discovery Grant (RGPIN-2016-04890) to J.D., a NSERC Grant (RGPIN 238338), a Canadian Institutes of Health Research Grant (PJT-153447), and a National Institute of Child Health and Human Development Grant (R01 HD075740) to P.L.G.

## Notes

**Conflict of interest** The authors declare no conflict of interest.

### Competing Interest Statement

The authors have declared no competing interest.

