## Appendix 1 for "The role of feedback in the production of skilled finger sequences"

**Subj:** **Study: Sequence Integration 4**

<u>Questionnaire about Experience:</u>

Did you notice anything during the experiment?

We manipulated an aspect of the task during the experiment what was it?

Which of these manipulations did we implement (chose any that apply)?

- ☐ Change the frequency of the tones that were presented when a key was pressed
- ☐ Delay the feedback of a press
- ☐ Provide false feedback on a press (if you were correct it would show as incorrect)
- ☐ Interleave the 3 trained sequences with random sequences
- ☐ Change the frequency of the vibration when a key was pressed
- ☐ Advance the feedback of a press
- ☐ Switched a single press within the sequence (switch which number is presented)
- ☐ Randomize the points you received for each trial rather than making them dependent on performance
- ☐ Omit the feedback of a press
- ☐ Give you false feedback regarding your average speed at the end of a block (higher or lower than you actually were)
